# Biodiversity State Indicator: Integrating taxonomy, functionality and species interactions

**DOI:** 10.1101/2025.05.23.655681

**Authors:** Lars O. Mortensen, Ole B. Brodnicke, Cecilie B. Devantier, Sofie Ferreira, Xin Huei Wong, Verena Schrameyer

**Affiliations:** DHI A/S, Section for Offshore wind - Environment, Hørsholm, Denmark; DHI A/S, Section for Marine & Coastal – Environmental Solutions, Hørsholm, Denmark

**Keywords:** biodiversity index, reporting standards, communication, GRI, Global Reporting Initiative, EU Taxonomy, TNFD

## Abstract

The biodiversity crisis is a pressing global issue, mainly driven by climate change and anthropogenic pressures. Efforts to mitigate climate change have had limited success, emphasizing the need to address local stressors to prevent biodiversity collapse. However, measuring biodiversity remains complex due to its multifaceted nature, including taxonomic, functional, and interaction dimensions.

This study introduces the Biodiversity State Indicator (BSI), an integrated metric designed to assess biodiversity by combining classic taxonomic biodiversity with functional biodiversity and community interactions. The BSI calculates the state of biodiversity at a target site, compared to reference ecosystems, enabling the indicator to assess net-positive or net-negative biodiversity change. The indicator is validated through simulations and applied to North Sea data, demonstrating its sensitivity to species richness and its capacity to track biodiversity changes over time.

The BSI was shown to perform accordingly to expectation, showing no preference for a specific metric included in the calculations. Additionally, the BSI showed that while giving a clear and simple number to communicate and relate state of biodiversity and biodiversity changes, resolving the calculations into its constituent components can assist in resolving some of the internal ecosystem dynamics by levering changes in taxonomic, functional and interaction diversity with each other. In conclusion, the Biodiversity State Indicator can be a solid tool to assist in the net-positive strategy of marine stakeholders, developers and decision makers.

## 1 Introduction

The biodiversity crisis is increasingly recognized as a pressing global issue (Singh, 2002; Isbell et al., 2023). Species declines are closely linked to climate change and intensified anthropogenic pressures, which are anticipated to drive large-scale ecosystem transformations in the coming decades (Meyer et al., 2022). In marine environments, all 29 coral reef World Heritage sites are projected to undergo significant deterioration, leading to reduced ecological functioning and a consequent decline in the quantity and quality of ecosystem services provided (Heron et al., 2017). Efforts to mitigate climate change and restrict global warming to 1.5°C have largely been unsuccessful (Matthews and Wynes, 2022), yet research (Brown et al., 2013) suggests that addressing local stressors is crucial for mitigating biodiversity collapses.

International frameworks such as the Convention on Biological Diversity (Secretariat of the Convention on Biological Diversity, 2020), the Global Reporting Initiative (Global Reporting Institute, 2024), and the European Union’s Green Deal (European Commission, 2019) have been pivotal in establishing guidelines for companies and governmental institutions to report on and engage with biodiversity. These initiatives promote sustainable practices by integrating biodiversity considerations into decision-making processes, emphasizing the critical role of biological diversity in maintaining ecological stability and human well-being.

Despite these efforts, a significant challenge remains: the absence of specific methodologies for comprehensive biodiversity measurement. Biodiversity is an intricate and intangible concept encompassing both taxonomic diversity - the variety of species, and functional diversity - the roles species play within ecosystems (Cardinale et al., 2012). Current frameworks often lack detailed guidance on quantifying these biodiversity measurements, resulting in inconsistencies in reporting and assessment. Moreover, the complexity of biodiversity is difficult to capture using traditional indicators, which typically focus on isolated either taxonomic or functional diversity. This limitation underscores the urgent need for integrated indicators that can combine diversity perspectives to provide a holistic and data-driven quantification of biodiversity. Integrated indicators are essential for accurately monitoring biodiversity changes, evaluating the effectiveness of conservation efforts, and guiding informed decision-making (Mace et al., 2018).

To address these needs, we introduce the Biodiversity State Indicator (BSI) that builds on an original Standardized Biodiversity Index (SBI) (Rey Benayas and De La Montaña, 2003; Assunção– Albuquerque et al., 2012; Chiatante et al., 2021). The novel composite approach is designed to benchmark the biodiversity state of a target ecosystem, by integrating taxonomic and functional diversity, along with interaction between species and functional traits. The BSI compares the biodiversity of a target site with a reference site, offering insights into the degree of biodiversity change and its potential ecological consequences. Thus, the aim of this paper is to demonstrate the performance of BSI to benchmark the biodiversity of a range of simulations. Furthermore, after validation process, we apply BSI to an empirical data set to demonstrate the usage of BSI in biodiversity communication and monitoring. Lastly, we demonstrate the applicability of BSI in predictive modelling.

Adopting integrated biodiversity indicators allows companies and governments to better align with international frameworks and contribute more effectively to global biodiversity conservation goals. These indicators bridge the gap between policy aspirations and practical implementation, ensuring that efforts to protect biodiversity are grounded in comprehensive and reliable data.

## 2 Methodology

In the current study we integrated taxonomic (Supplementary: Table 1), functional (Supplementary: Table 2), and interaction (Supplementary: Table 3) diversity into a biodiversity quantification benchmark because each dimension offers unique insights into ecosystem complexity and resilience: Taxonomic Diversity: Provides a foundational understanding of species richness and composition, essential for assessing overall biodiversity. It helps identify areas with high species diversity and potential conservation value.

Functional Diversity: Captures the range of functional traits and ecological roles that species have within an ecosystem. This dimension is vital for understanding how ecosystems function and deliver services such as nutrient cycling, habitat formation, and resilience to environmental changes. Functional diversity links biodiversity to ecosystem service capacity, offering insights into how ecosystems support human well-being.

Interaction Diversity: Focuses on the relationships and interdependencies among species, crucial for maintaining ecosystem stability and dynamics. Understanding these interactions, including trophic links and species networks, can reveal how ecosystems respond to disturbances and how energy and resources flow through ecological communities.

By integrating these three dimensions of biodiversity, we obtain a holistic view of ecosystem health and complexity, which is essential for effective conservation and management strategies. This comprehensive approach allows for a more nuanced understanding of biodiversity patterns and processes, facilitating the identification of key species and interactions that sustain ecosystem services and resilience.

### 2.1 Calculations of the Biodiversity State Indicator (BSI) capturing biodiversity change

The BSI is designed as a single value benchmark, indicating if the target ecosystem has a higher or lower biodiversity than a reference ecosystem, which can be a geographical or temporal reference site/scenario. This was done to support frameworks that emphasize net-positive, net gain or net-zero biodiversity. However, as biodiversity is an abstraction of a multidimensional concept (Purvis and Hector, 2000), we here define biodiversity as an integrated concept, comprising of taxonomic diversity, functional diversity and the degree of possible interactions between ecosystem components (Connectivity, I-diversity). Generally, the individual diversity types are calculated for both the target and reference ecosystem and the target ecosystem is then benchmarked against the reference ecosystem. A more detailed description of the calculation is described in the following section.

#### Definition of target and reference ecosystem

To accurately assess net changes in biodiversity, it is essential to establish a baseline or reference state of biodiversity. This reference state can be defined either temporally, such as the biodiversity status prior to a construction project or environmental impact, or spatially, representing a known desired biodiversity condition in a specific area. Establishing this baseline is a critical step in biodiversity assessment, as it provides a benchmark for measuring deviations and changes over time or across different locations.

Data collection should encompass the species present and their respective abundances in both the target and reference sites. It is crucial to ensure that data collection efforts are consistent and comparable between these sites. This requires applying equal sampling effort and methodology to both locations to avoid biases that could skew the results. The data collected should cover the same ecological parameters and habitat types at both sites. For instance, if data at one site focuses solely on benthic communities, the data collected at the other site should be limited to benthic communities as well. This consistency in data resolution and scope is vital for ensuring that any observed differences in biodiversity are due to actual ecological changes rather than discrepancies in data collection methods. The resolution of data, or the level of detail captured in the dataset, must be equivalent across sites to ensure accurate comparisons. This includes factors such as the spatial and temporal scales of sampling, the taxonomic resolution (i.e., how finely species are identified), and the ecological functions or interactions being measured. High-resolution data allow for more precise analyses and interpretations of biodiversity changes, enabling researchers to detect subtle variations that might otherwise go unnoticed.

#### Calculating α-, β- and I-diversity

Initially, standard diversity indices are calculated for target and reference site. Taxonomic diversity indices currently included in the BSI are listed in Table 1, Functional diversity indices in Table 2 and Interaction diversity indices in Table 3.

**Table 1.** Indices used for quantification of taxonomic diversity.

| Type | Index | Description |
| --- | --- | --- |
| Richness | Margalefs d | Weighting the number of species with the number of individuals. |
| Evenness | Simpsons index | A measure of diversity that takes the number of individuals within each species identified into account. (Simpson, 1949) |
| Evenness | Pielou index | Shannon index in relation to individual species abundance (Pielou, 1966) |
| Uniqueness | Invasiveness | Number of invasive species identified in the samples according to local and global data bases |
| Uniqueness | Rareness | Average IUCN red list score across species identified along with a weighted average between number of individuals of each species and red list score. |

**Table 2.** Indices used for quantification of functional diversity.

| Type | Index | Description |
| --- | --- | --- |
| Other | AMBI index | Assessment of seabed community quality. Can function as an index of sensitivity traits (Golumbeanu et al., 2014) |
| Other | Community weighted mean (CWN) | Weighted mean trait size in the community. Can be used to compare communities. |
| Other | Functional rarity | Use MPD or MNTD on individual species |
| Richness | Trait range | An indication of the variability within each trait. Combining the variability of each trait into a convex hull to account for functional diversity of the traits. (Cornwell et al., 2006) |
| Richness | Variance | Variance in traits ranges formulated as $\sum_{i=1}^N p_i (x_i - CWM)^2$, where CWM is the community weighted mean, pi is the proportional species abundance and xi is the trait value of species i. |
| Richness | Number of functional groups | As with taxonomic diversity, the pure number of functional groups represents a measure of diversity, without considering the abundance within each group. |
| Evenness | Regularity | Indicator of how regular species are distributed along trait ranges. Can be measured using indices like FEve (Villéger et al., 2008) or Mean Neighbor Taxon Distance (MNTD)(Webb et al., 2002) |
| Divergence | Mean Dissimilarity | This measures the mean dissimilarity in traits between species and has several approaches to calculate, such as the Mean Pairwise Dissimilarity (MPD)(Weiher et al., 1995). Rao Quadratic Entropy (de Bello et al., 2016) or Functional Dispersion(Laliberte and Legendre, 2010). |

**Table 3.** The indices used for quantification of interaction diversity.

| Index | Description |
| --- | --- |
| Network density | The ratio of observed number of connections and the maximum possible connections. |
| Diameter of network | Shortest distance between the most distant nodes in the network |
| Average path length | The average distance between each node in the network. |
| Trophic guild ratio | A ratio between all the trophic guilds across the community surveyed. |

Taxonomic diversity, defined as the mean and variance of the number of species within a target ecosystem (Neeson et al., 2013), has a long-standing tradition in ecology and has historically served as the primary measure of biodiversity (Warwick and Clarke, 1998). Over the decades, various indices have been developed to describe biodiversity as a function of the number of species and individuals, with each index capturing different aspects of taxonomic diversity. To provide a comprehensive view of taxonomic biodiversity, we calculate several indices, each highlighting distinct aspects of species richness and abundance (Teixeira et al., 2016).

We also incorporate indices of functional diversity. For this purpose, data from the North Sea were integrated with traits defined in the CEFAS trait database (Clare et al., 2022), which details ten key biological traits of marine benthic invertebrates surveyed in Northwest Europe. This grouping allows for an analysis of functional ecological traits, providing a more comprehensive description of target ecosystem functioning and linking it to ecosystem service capacity.

The final dimension of biodiversity considered is the interactions between identified taxa. Interaction diversity is quantified through indices based on abundance, taxonomic identity, and trophic relationships. Three of these indices derive from interconnectivity among species, assessed through species networks constructed using species co-occurrence data. These networks were calculated using the Igraph package (Csárdi et al., 2023) in R (version 4.3.1, “Beagle scouts”). For each station and reference site, we calculated network density, diameter, and average path length. Additionally, the trophic guild ratio, defined as the mean ratio among all trophic guilds at a station, was calculated.

#### Standardization and aggregation into BSI

Finally, the indices are standardized and benchmarked against the reference site, resulting in an aggregated benchmark score, called the BSI. First, each index from the target site is standardized against a reference scenario using the equation:

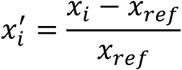

Where 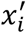 is the standardized index, *x*_*i*_ is the station index, *x*_*ref*_ is the reference station index. Standardization allowed the indices to be dimensionless and on the same scale for comparison. The standardized values of the indices act as a scoring for each station, with the reference stations set to 0. The index scores of all indices in the diversity group are then averaged for a diversity score at the station.

Next, the average across standardized indices within each compartment (taxonomic, functional and interactions) were calculated, yielding a partial BSI for taxonomic diversity, functional diversity and interactions. Lastly, the partial BSI’s were averaged to a joint Biodiversity State Indicator (BSI), which is the final benchmark, indicating if the general biodiversity is higher or lower at the target site, than the appointed reference site.

### 2.2 Validation

#### Study site

This study was conducted using a hybrid approach, combining simulation models with real-world data from the North Sea. The simulation aspect allowed for controlled testing of specific variables, while the empirical data provided insights from actual environmental conditions. The North Sea, a shallow continental shelf sea located between the UK and mainland Europe, supports a rich and diverse ecosystem. It is home to various fish species, marine mammals, and seabirds, and plays a crucial role in regional fisheries and biodiversity. The area is influenced by strong tides and currents, which shape its unique biological and ecological characteristics.

#### Simulation data: Number of species

Using the species list provided by Clare et al. (2022), we generated 1 000 sets of genus lists, each containing a random number of genera. From these, we selected one set comprising 66 genera, based on our experience with other sampling programs in the North Sea, where this number is considered representative of a typical sampling site. For each genus in this set, a random number of individuals was assigned, ensuring that the total number of individuals at each site summed to 20 000. The BSI algorithm was then applied to each dataset, and the resulting BSI scores and associated metrics were analyzed to evaluate the impact of varying species numbers on the BSI score.

#### Simulation data: Number of individuals

To test the effects of varying individual counts on the BSI score, we generated 1 000 datasets with the same genus composition as the reference site selected in the previous step, but with different numbers of individuals assigned to each genus. Each dataset was then analyzed using the BSI algorithm, and the resulting BSI scores and underlying metrics were examined.

#### Simulation data: Number and type of traits

To assess the potential influence of the number and types of traits on the BSI score, we developed a trial dataset based on the reference set from the initial step. We first calculated the BSI using the complete set of traits described by Clare et al. (2022). Subsequently, we re-ran the analysis, systematically varying the number and types of traits included. This process was repeated for all possible trait combinations.

#### Real world data

The BSI algorithm was also applied to seabed infauna data from the North Sea, which is part of routine seabed monitoring conducted around oil and gas platforms within the Danish Exclusive Economic Zone (EEZ). This dataset comprises sediment core samples collected from 14 stations surrounding a single platform during the years 2006, 2009, 2015, and 2018. Each station was sampled using a Haps core sampler, with seven subsamples taken per site. The subsamples were sieved through a 2 mm mesh and preserved in ethanol before being analyzed at the Danish Biological Laboratory (DBL). Taxonomic identification was performed to the lowest possible level, resulting in a list of taxa identified at each station. Density estimates were calculated based on the number of individuals found in the samples, divided by the sampled area (145 cm^2^). This species list was then used for further processing and analysis.

Species traits related to their ecological functions were assigned at the genus level, based on information from Clare et al. (2022). Where possible, identified species were aggregated to the genus level. A total of 10 traits were associated with the genus list. On average, nine taxa per year lacked trait information due to missing data.

In this test, the BSI was calculated using two types of reference sites. The first reference site was the standard reference site used in the monitoring program, representing a location with minimal anticipated anthropogenic impacts. The second reference site was a worst-case scenario, where all biodiversity indicators were set to their lowest possible values. This approach was employed to evaluate whether the BSI could effectively quantify biodiversity when compared to a baseline of zero biodiversity.

## 3 Results

### 3.1 Validation analyses

#### Sensitivity to number of species

There was a significant effect of the number of species on the overall BSI (linear regression, p < 0.01, Fig. 1A). This effect was derived from an effect of the number of species on both the taxonomic (linear regression, p < 0.01, Fig. 1D) and functional diversity (linear regression, p < 0.01, Fig. 1C). The interaction diversity seemed unaffected by the number of species.

**Figure 1.**
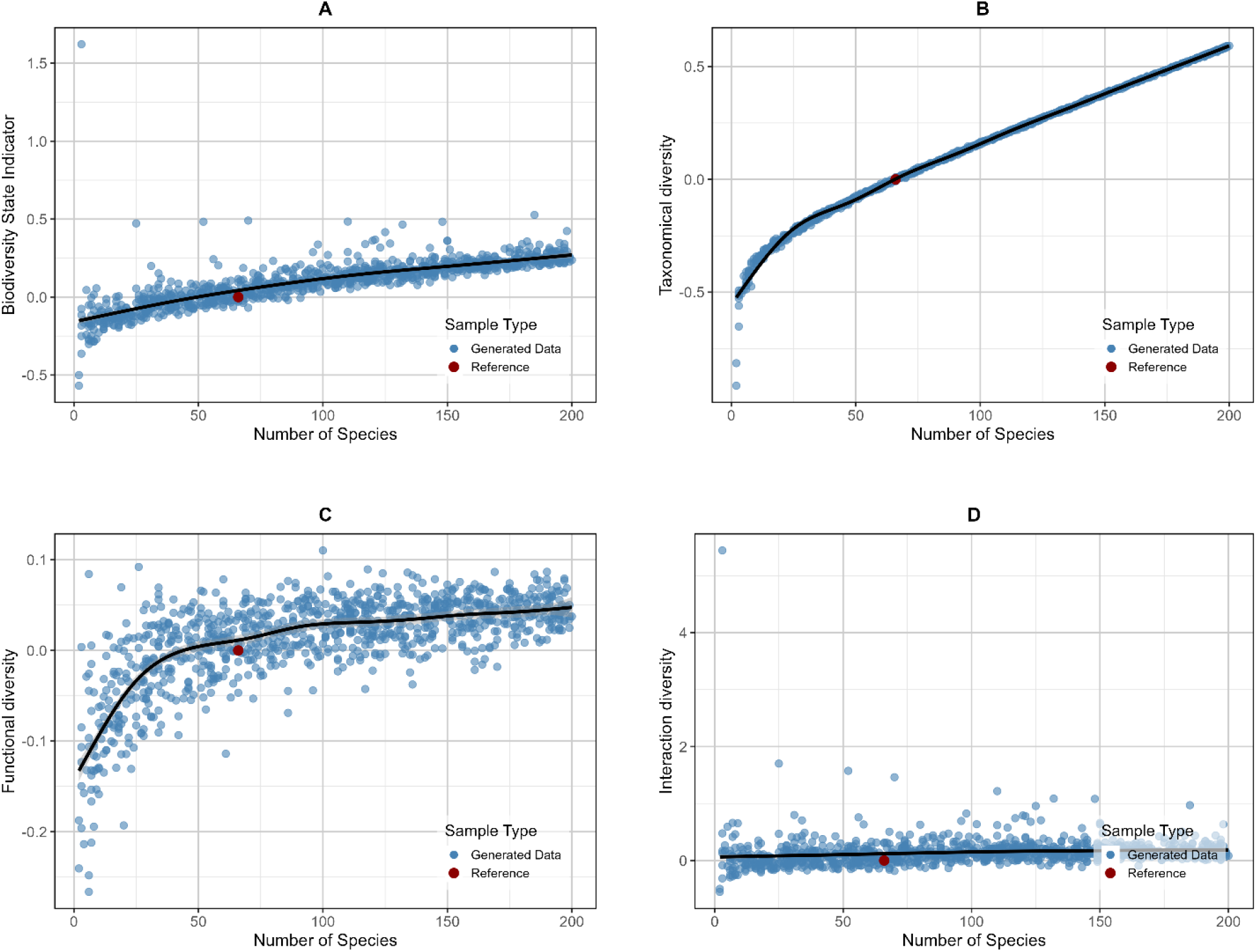
Relationship between the number of species in a sample and the BSI (A), average taxonomic diversity (B), average functional diversity (C) and average interaction diversity (D). Black line indicates a general additive model smoother to indicate trends.

#### Sensitivity to number of individuals

There was a significant decrease in BSI as a function of the number of individuals observed in the same set of species (Figure 2 A) (linear regression, p<0.01). The small change is because of individuals on the taxonomic diversity, where the Marglef’s d is very dependent on the number of individuals (Figure 2 B). Neither the functional nor interaction diversity was affected by the number of individuals (Figure 2 C & D).

**Figure 2.**
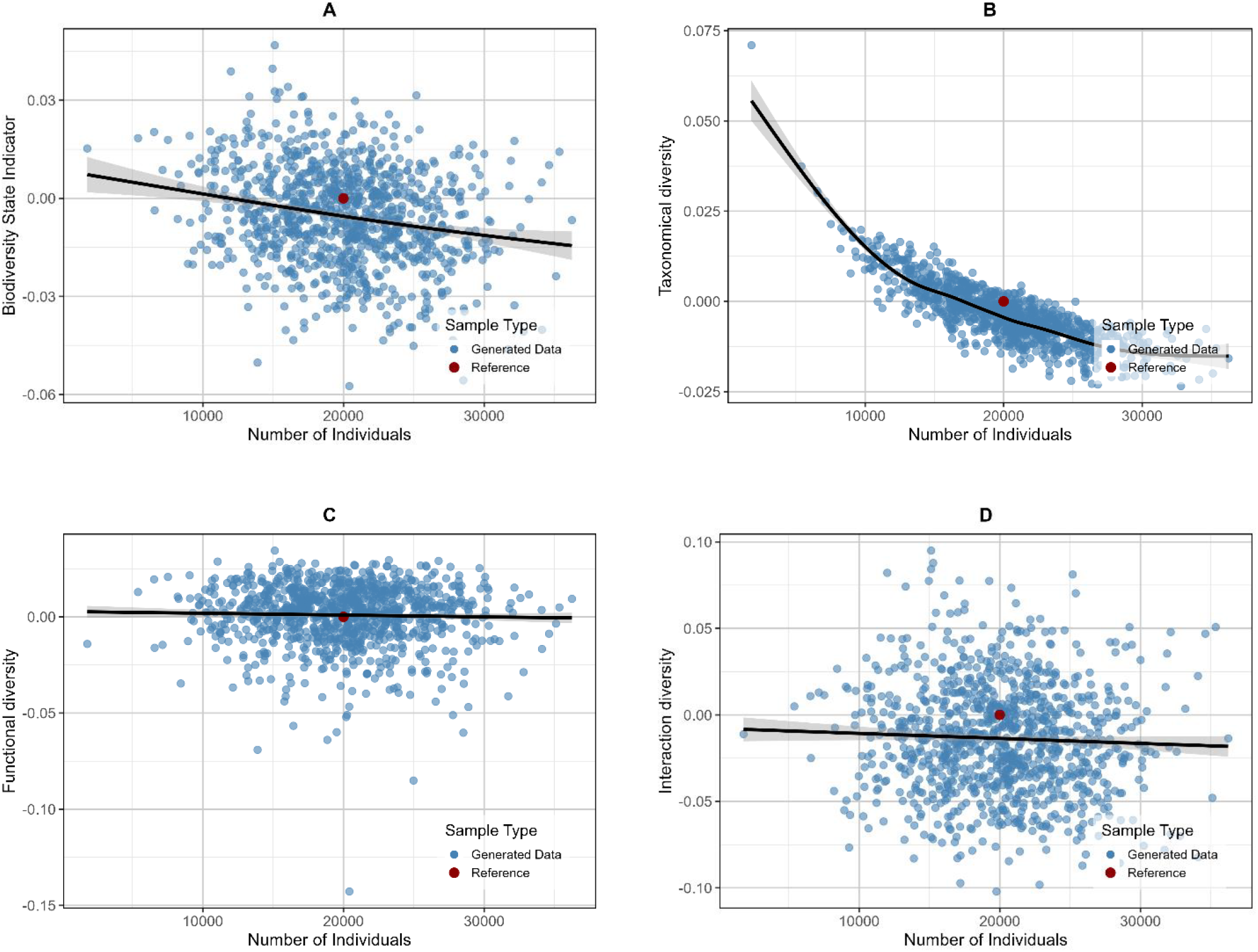
Relationship between the number of individuals in a sample and the BSI (A), average taxonomic diversity (B), average functional diversity (C) and average interaction diversity (D). Black line indicates a general additive model smoother to indicate trends.

**Figure 3.**
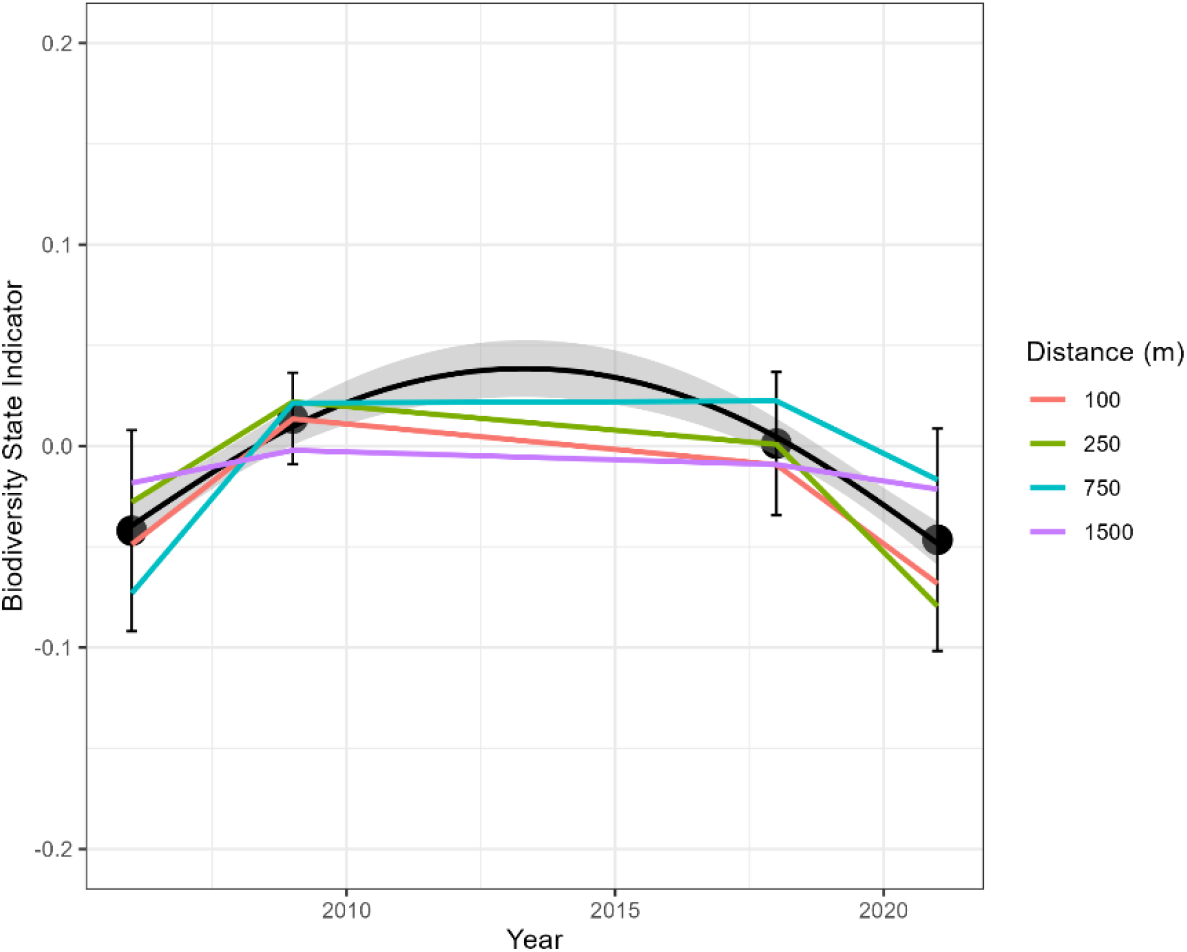
Development in the standardized BSI diversity from 2006 to 2021. Black line indicate gam smoother (BSI∼s(Year,k=3)), and colored lines indicate scores at stations at different distances to the platform. Each distance has 4 sample sites at each corner of the compass.

### 3.2 Real world data

The biological surveys conducted in the North Sea detected 139 taxa across all sampling years and stations. The number of total taxa detected was variable, ranging from 69 in 2009 to 78 in 2021. The taxa abundance was variable with 55.6 ± 243.8 SE individuals per 0.1 m^2^ across all stations and sampling years. The biological survey also includes the sampling at a regional reference site, located +20 km away from the platform. This site was selected as a reference, using the year 2006 to have a steady baseline measurement, to benchmark the platform samples.

#### *Aggregation into* BSI

The aggregated BSI score across all years and stations were calculated to be -0.01 (± 0.04 SD) indicating that the biodiversity overall around the platform was lower than on the reference station. However, resolving the BSI score showed that non-linear change in the score (GAM, p > 0.01).

The temporal trends in the BSI changed per index group (*Figure 4*). The taxonomic diversity had a significant change in time (linear regression, *x*^2^, p < 0.01, R^2^ = 0.57), while the functional diversity significantly decreased in time (linear regression, p<0.01, R^2^ = 0.40), although there where a small increase in 2009 followed by a decrease. There was also a significant decrease in the interaction diversity (linear regression, p < 0.01, R^2^ = 0.06).

**Figure 4.**
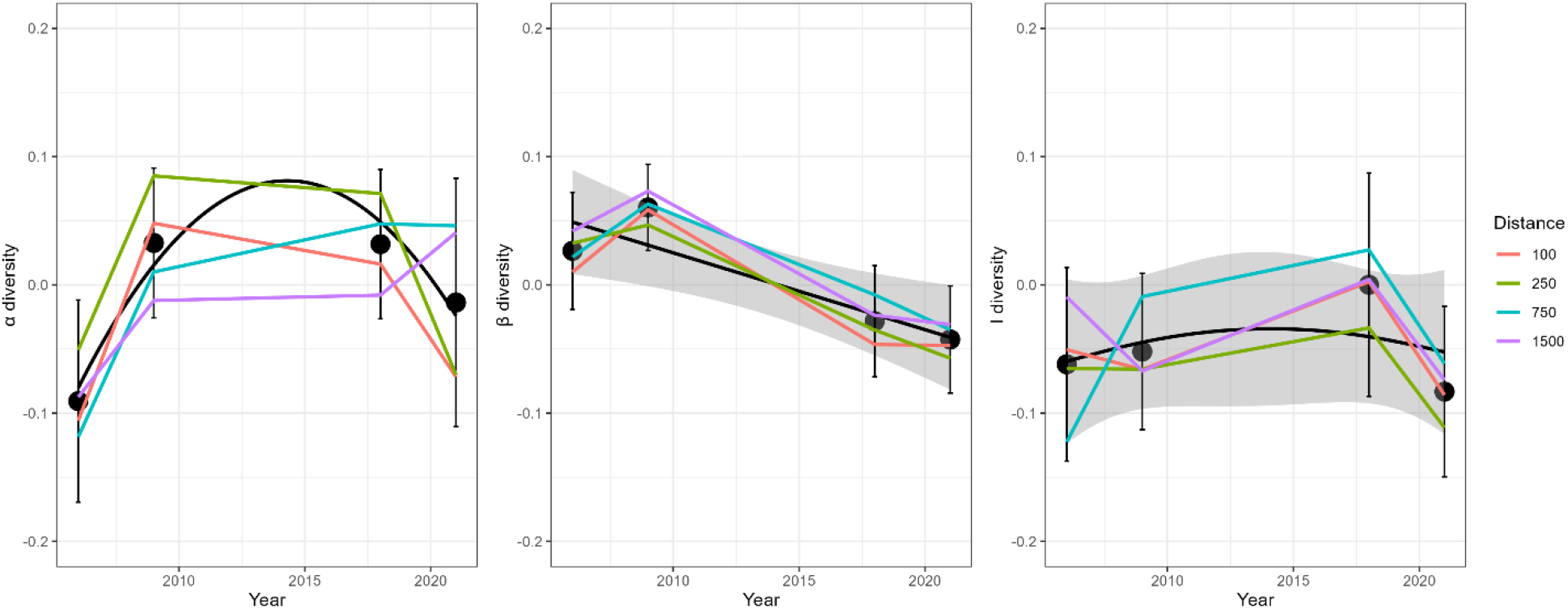
Development in the individual index groupings of the BSI diversity from 2006 to 2021. Black line indicate gam smoother (BSI∼s(Year,k=3)), and colored lines indicate scores at stations at different distances to the platform. Each distance has 4 sample sites at each corner of the compass.

**Figure 5.**
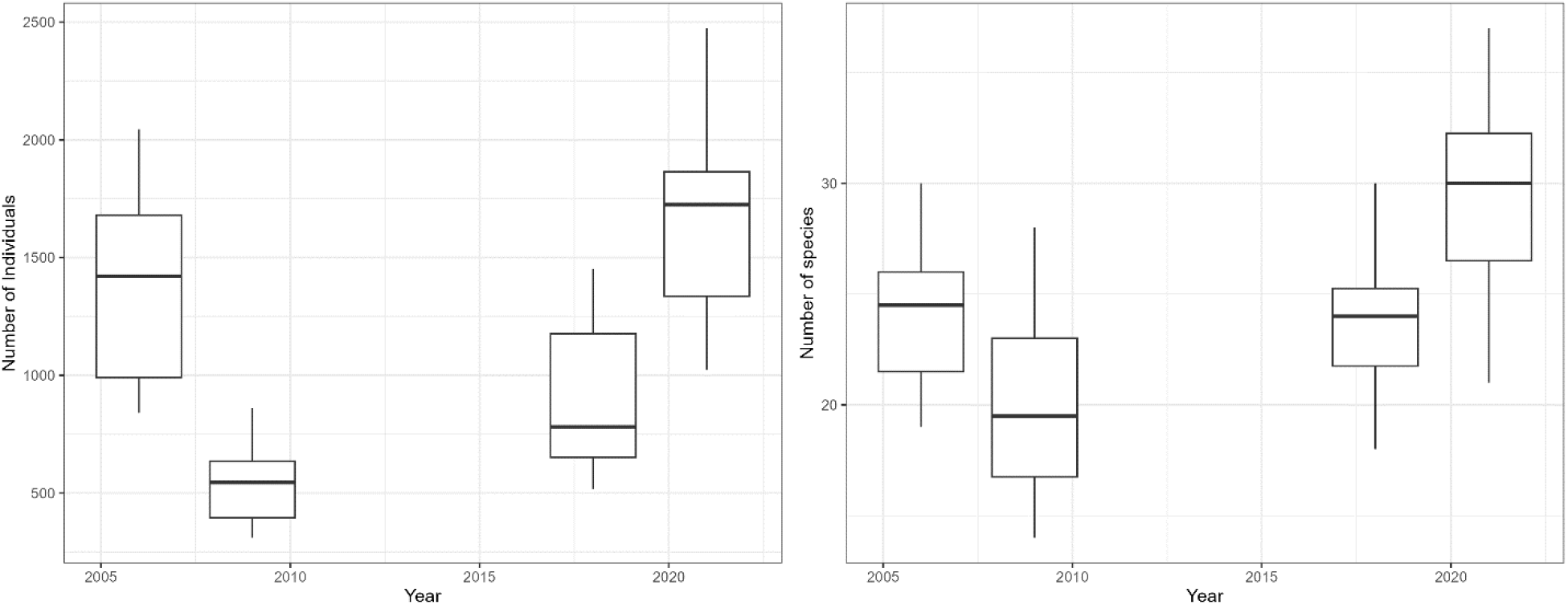
Average number of individual (left) and average number of species (right) found across all stations and years.

#### Detailing the Indicator

To detail the identified indices, all three were resolved to the individual indices they were based on and compared with the number of species and individuals found at the station.

Linking the overall BSI score to the number of species and individuals at the corresponding station showed a significant relationship between the two variables and the BSI score, with the score increasing with the number of species at the station, while decreasing with the number of individuals (Figure 6).

**Figure 6.**
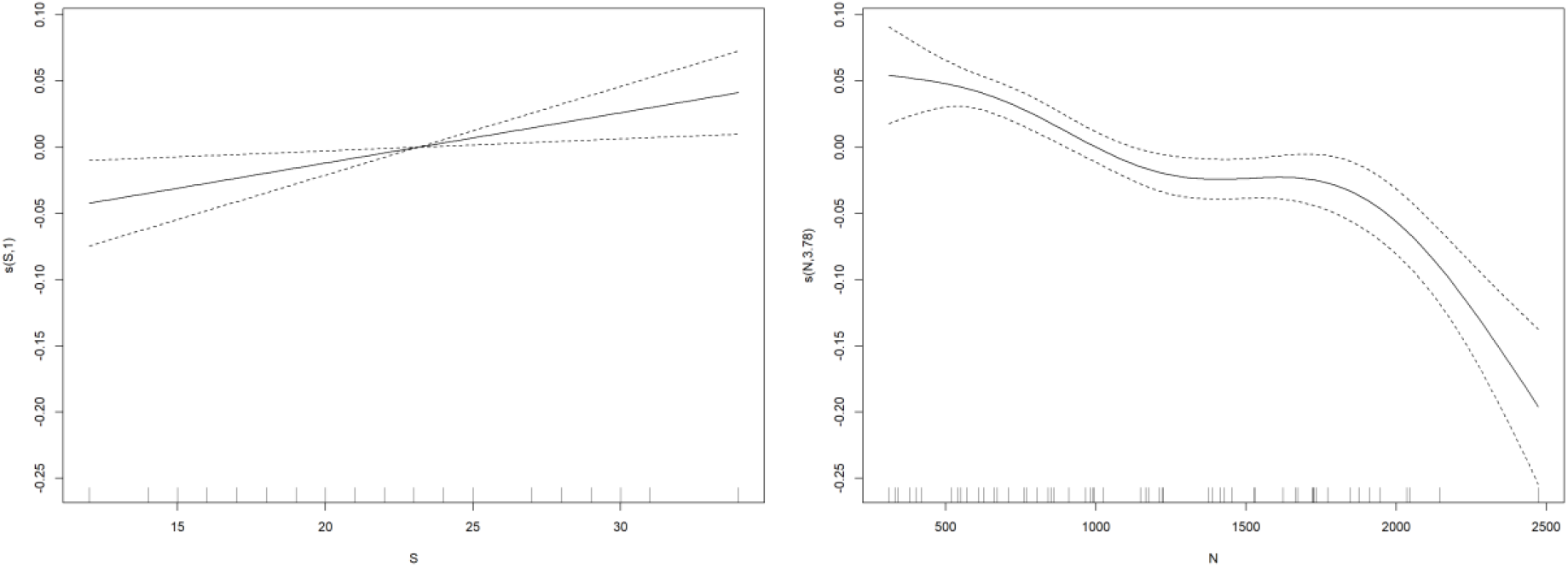
Results from GAM models linking the BSI to the number of species and individuals observed at each station.

Resolving the BSI into the index groups and analyzing the relationship between the calculated average taxonomic, functional and interaction diversity and the number of species and individuals (Figure 7) showed that the taxonomic diversity increased with the number of species, while it decreased with number of individuals (GAM, p < 0.01, R^2^ = 0.62)

**Figure 7.**
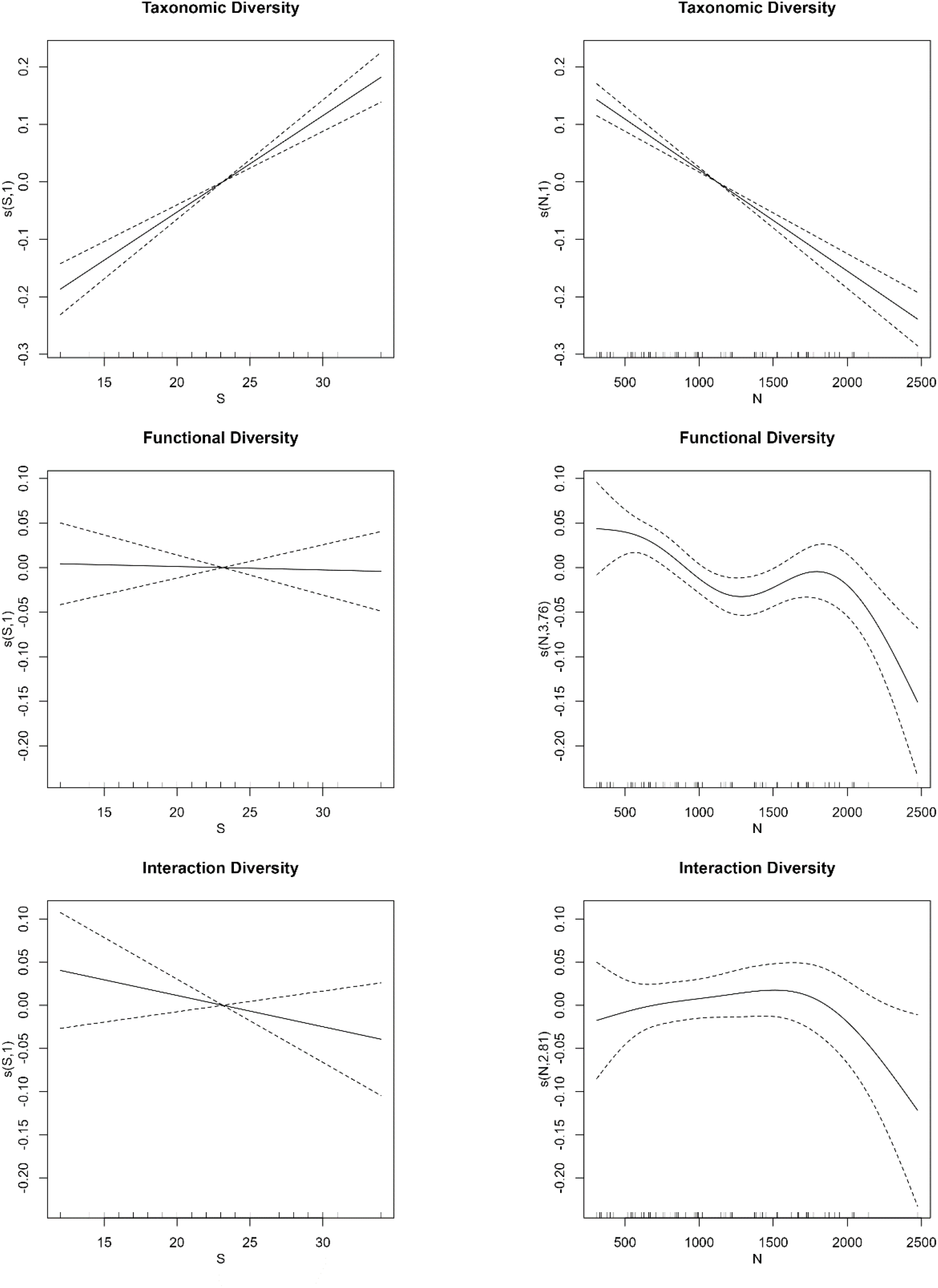
Results from GAM models linking the taxonomic, functional and interaction diversity to the number of species and individuals observed at each station.

Based on the detected number of species and individuals at each platform station (**Error! Reference s ource not found**.), we also analyzed the change in each individual diversity index (*Figure 8*). There was a significant difference within the taxonomic indices, where the Margalef’s d were inversely proportional to the Simpson, Shannon and Pielou index. Similarly, the functional diversity indices were also not uniform across the years, with functional richness being distinct from the others, while the RaoQ and functional dissimilarity had similar curves. Also, the interaction diversity curves were different from each other.

**Figure 8.**
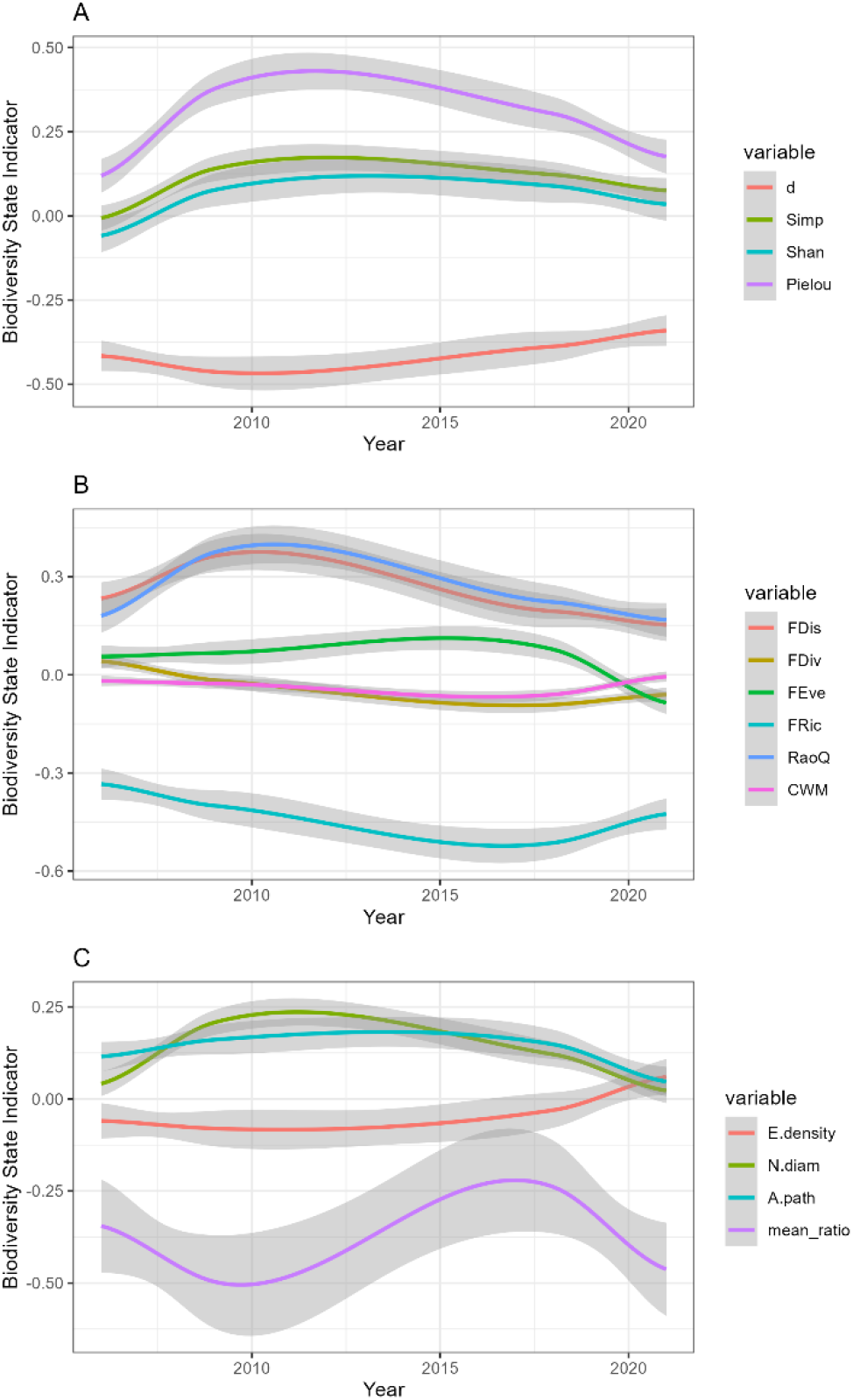
LOESS smoothing functions for individual diversity indices for α (A), β(B) and I(C) diversity across time for all stations.

## Discussion and Conclusion

### Sensitivity of the Biodiversity State Indicator (BSI)

The study’s simulations demonstrated that the BSI is sensitive to variations in species richness, with a significant positive correlation between the number of species and the overall BSI score. This finding suggests that taxonomic diversity heavily influences the BSI, reinforcing the notion that species richness remains a crucial component of biodiversity assessment. The positive relationship indicates that ecosystems with a higher number of species are likely to exhibit higher biodiversity scores, reflecting greater ecological complexity and potential resilience.

Interestingly, the study also found that while the number of species significantly affected both taxonomic and functional diversity, interaction diversity remained largely unaffected by species richness. This result highlights a critical aspect of biodiversity: the stability of species interactions might be less sensitive to changes in species numbers compared to other biodiversity dimensions. This suggests that even in ecosystems experiencing species loss, the network of interactions might persist, although the ecological roles and services provided by the ecosystem could be compromised.

Regarding species abundance, the study revealed a significant but relatively minor effect on the BSI score. The relationship between species abundance and taxonomic diversity, as measured by Margalef’s d index, indicated that while more individuals contribute to higher diversity scores, this effect plateaus, suggesting diminishing returns with increasing abundance. This nuance is important as it underscores that mere increases in individual numbers do not necessarily translate into proportionally higher biodiversity, especially if those numbers are skewed towards a few dominant species.

### Real-World Application of the BSI to North Sea Seabed Data

Applying the BSI to real-world data from the North Sea provided valuable insights into its practical application. The analysis of seabed infauna around oil and gas platforms revealed a nuanced picture of biodiversity changes over time. The overall BSI score of -0.01 indicates a slight decline in biodiversity at the platform sites compared to the reference site, suggesting that industrial activities may have had a subtle but persistent impact on local ecosystems.

When disaggregated, the results showed divergent trends among the different biodiversity components. Taxonomic diversity around the platforms increased over time, potentially indicating either an adaptation to the altered environment or a shift in species composition favoring more tolerant species. This increase could also reflect the introduction of new species or the recovery of certain taxa after initial disturbances, which is a critical consideration for monitoring programs aiming to track long-term ecosystem health.

Conversely, the study found a significant decline in both functional and interaction diversity over time. The decrease in functional diversity suggests that while the number of species might have increased, the range of ecological roles and functions they perform has narrowed. This reduction in functional diversity can have profound implications for ecosystem resilience, as fewer functional roles may lead to diminished ecosystem services, such as nutrient cycling and habitat formation, which are essential for maintaining ecological balance.

The observed decline in interaction diversity further reinforces concerns about the long-term stability of these ecosystems. A reduction in the complexity of species interactions could lead to less resilient ecological networks, making the ecosystem more vulnerable to additional stressors, such as climate change or further anthropogenic impacts. This decline is particularly concerning in the context of marine ecosystems, where trophic interactions play a vital role in maintaining the structure and function of the community.

### Implications for Biodiversity Monitoring and Management

The study’s findings highlight the value of the BSI as a tool for biodiversity assessment, particularly in environments subject to anthropogenic pressures. The ability of the BSI to integrate multiple dimensions of biodiversity provides a comprehensive measure that can reveal underlying changes not captured by traditional methods focusing solely on species counts. The differential trends observed among taxonomic, functional, and interaction diversity emphasize the importance of a multidimensional approach to biodiversity monitoring.

For management and conservation efforts, these results suggest that while species richness is a valuable indicator, it should not be the sole focus. The declines in functional and interaction diversity indicate potential losses in ecosystem resilience and function, which could lead to more significant and possibly irreversible changes in the future. Therefore, monitoring programs should incorporate integrated measures like the BSI to capture these subtle yet critical changes in ecosystem dynamics.

Moreover, the study demonstrates that the BSI can be effectively applied to real-world data, providing actionable insights into biodiversity changes over time. This capability is particularly relevant for regulatory and conservation frameworks, which often require robust, evidence-based assessments to guide policy decisions and management strategies. By providing a more detailed and nuanced picture of biodiversity, the BSI can help ensure that conservation efforts are both targeted and effective, ultimately contributing to the long-term sustainability of ecosystems.

In conclusion, the study underscores the importance of using integrated biodiversity indicators like the BSI to monitor and manage biodiversity, especially in regions experiencing significant anthropogenic impacts. The results obtained through data validation and real-world data applications highlight the utility of the BSI in capturing complex ecological changes, making it a valuable tool for researchers, policymakers, and conservation practitioners.

## 5 Acknowledgments

We would like to thank TotalEnergies for the use of seabed infauna data, derived from one of their North Sea platforms. We also thank all anonymous reviewers for constructive discussion and input.

